# Correlation between DNA double strand breaks and cell death in peripheral blood lymphocytes from breast cancer patients

**DOI:** 10.1101/2024.12.23.630130

**Authors:** Ángela Solana-Peña, Monica Pujol-Canadell, Juan-Sebastián López, Miquel Macià, Evelyn Martínez Pérez, Isabel Linares, Milica Stefanovic, Héctor Pérez-Montero, Javier González-Viguera, Marina Arangüena Peñacoba, Montse Ventura, Gisela de Miguel-Garcia, Ferran Guedea, Nadina Erill, Victor González-Rumayor, Gemma Armengol, Joan Francesc Barquinero

**Author notes:** Corresponding author: Joan Francesc Barquinero Unit of Biological Anthropology, Department of Animal Biology, Plant Biology and Ecology, Autonomous University of Barcelona, 08193 Cerdanyola del Vallès, Spain.

## Abstract

Radiotherapy is an effective treatment to fight cancer. However, it not only affects cancer cells but also healthy tissues, causing side effects. Different factors can influence the appearance of radiotoxicity, like total dose administered or patient individual characteristics, such as genetic variability. Several biomarkers have been proposed to predict radiotoxicity, especially those based on apoptosis or DNA damage, for example γ-H2AX, which correlates with DNA double strand breaks. Our purpose is to analyze how apoptosis and γ-H2AX correlate to each other and to link these results with selected SNPs associated with apoptosis. Blood samples from 60 breast cancer patients in remission were recruited. After mononucleated cells isolation, samples were irradiated. Then, we assessed induction and kinetics of disappearance of γ-H2AX at different times after 2-Gy irradiation and apoptosis induced 24 and 48 h after 8-Gy irradiation. A negative correlation was observed between basal and residual γ-H2AX and apoptosis at 48 h post-irradiation. This result supports previous studies with cancer patients showing a negative correlation between these two biomarkers. Considering the high variability of radio-induced apoptosis, we performed a genotyping study. Two SNPs located at *TP53* and *FAS* genes were associated with apoptosis. Overall, our results indicate that individuals with less efficiency in removing damaged cells, probably due to genetic polymorphisms, presented more basal and residual levels of DNA damage.

## 1. Introduction

Radiation therapy (RT) is a common and effective treatment for several types of cancer thanks to the ability of high-energy radiation to kill cancer cells or inhibit their growth. According to the American Cancer Society, more than 50% of patients with cancer receive RT [1]. Although it can be administrated concomitantly with other treatments, such as chemotherapy or immunotherapy, RT is applied as monotherapy in about 15% of patients with solid cancer [2]. However, RT not only affects cancer cells but also healthy tissues or organs, which can develop adverse side effects, like erythema, ulcers and fibrosis [3]. Currently, technologic advances in RT, like stereotactic RT or intensity modulated RT, have improved dose conformation, delivering the maximum dose to the tumour and diminishing the exposure of healthy tissue and hence avoiding healthy tissue damage [3–5]. Despite this, it is accepted that between 5 and 10% of patients treated with RT will present adverse tissue reactions [6,7]. In accordance with the Hsu hypothesis [8], it is assumed that variation of healthy tissue reactions follows a Gaussian distribution, in which individuals affected by rare mendelian diseases that confer radiosensitivity, such as ataxia telangiectasia, Nijmegen syndrome, Fanconi’s anaemia and Bloom syndrome, represent the most sensitive cases. However, the incidence of these diseases does not explain the assumed percentage of 5 to 10%. Several factors can influence the observed variation, some of them are related to treatment itself (total dose administrated, fractionation dose, irradiated volume and irradiated body part), additional treatments (concomitant chemotherapy and/or surgery), or patient individual characteristics (age, sex and co-morbid conditions) [9,10]. But the characteristics mentioned above only explain about 20% of the variation and the remaining variation is likely due to genetic factors [11,12].

Due to the increasing number of cancer patients and therefore, of patients who will receive RT in the future, there is an interest in the search of biomarkers capable of predicting the toxicity resulting from RT. Currently, the most studied biomarkers are those based on DNA damage and on cell death in response to radiation, as well as germline genetic variants or radio-induced changes in gene expression [6,13–15]. In this sense, one of the earliest DNA damage detection events is the phosphorylation of the histone H2AX (γ-H2AX) at the DNA sites with double-strand breaks (DSB) [16,17]. This analysis of γ-H2AX phosphorylation has been suggested as biomarker of radiotoxicity in several studies. Some of these studies have described that the DSB repair capacity is diminished in cancer patients with RT toxicity [18,19]. Other studies have proposed the residual post-irradiation γ-H2AX foci as a radiosensitivity predictor [16,20–22].

Another group of well-established DNA damage biomarkers are based on cytogenetic techniques [23]. Some studies have seen an increase of radio-induced chromosomal aberrations in patients with normal tissue toxicity effects after RT [24–27]. Moreover, by using the G2 assay, several authors have found that the lower ability to arrest cell cycle damaged cells is related to radiosensitivity in breast, head and neck, colorectal and prostate cancer, as well as in some types of paediatric cancer [12,28,29] Furthermore, evaluating chromosomal aberrations by premature chromosome condensation techniques, some studies have shown an increase in extra chromosomal fragments in patients that suffer RT side effects [30–32]. Finally, some previous works have suggested that micronuclei, which are formed when a broken chromosome fragment remains outside the nucleus, are a good marker to detect patients with increased radiosensitivity [33,34].

The ability of cells to enter in programmed cell death or apoptosis is another possible biomarker of radiation sensitivity [35]. It has been proposed that *in vitro* radio-induced apoptosis of cytotoxic T lymphocytes may help to predict late toxicity induced by RT. Some studies show that patients with side effects after RT present lower levels of apoptosis than patients without them [36–39]. Variability in apoptosis levels depends on several factors, like the influence of genetic polymorphisms and their interactive network [40,41].

Genetic variations are assessed in radiogenomic studies with the aim of proposing them as biomarkers capable of predicting which individuals may develop adverse effects after RT. There are different types of studies, such as those in which differences in gene expression are studied [42] but also those in which different single nucleotide polymorphisms (SNP) are the target of study [43]. In the latter, the reported genes participate in different pathways that may affect the response to radiation, including apoptosis (e.g. *TP53*), cell proliferation (e.g. CDKs), cell cycle regulation (e.g. *ATM* or *CHK2*) or DNA repair (e.g. *XRCC4*). Additionally, genes associated with the response to oxidative stress, as well as those related to inflammatory and pro-fibrotic cytokines, have also been related to RT [44–46]. However, these findings have not been replicated in another cohort [47].

Notably, most of the above-mentioned studies have analyzed the radiosensitivity biomarkers individually. Few of them have tested the correlation between markers and particularly with genetic variants. Considering all the evidence mentioned above, the objective of the present work is to study how apoptosis and a DNA damaged biomarker, such as γ-H2AX, correlate to each other in a group of breast cancer patients and to link these results with selected SNPs associated with apoptosis.

## 2. Materials and methods

### 2.1 Patients

The study comprised 60 females, aged between 43 and 73 years, who had been treated for breast cancer and who were recruited for the study between 3 and 15 years after therapy. All of them were in complete remission. Patients were recruited in the Radiation Oncology Department at Institut Català d’Oncologia, Barcelona, Spain. The recruitment period was from November-2020 to March-2023. All patients received RT, 40 of them with a boost treatment and the mean dose received was 57.31 ± 8.9 Gy. Out of these patients, 33 were also treated with chemotherapy, 50 with hormonotherapy and 7 with monoclonal anti-HER2 antibody. This study was approved by the Research Ethical Committee of the Bellvitge Hospital and Institut Català d’Oncologia (PRED-RAD, reference PR044/20). All patients signed a written informed consent before blood collection and research methods followed Good Clinical Practice (GCP) guidelines and were performed in accordance with the Declaration of Helsinki. Clinical data needed in this study was only available by clinicians. Anonymized information was available from September-2024 to November-20214 for the other authors for research purposes.

### 2.2 Peripheral blood samples obtention

During a follow-up medical appointment, a peripheral blood sample of 27 ml was obtained by phlebotomy. Out of these ml, 18 ml of blood were collected in sodium heparin tubes (Vacutest, Arzergrande, Italy), 9 ml of blood were placed in EDTA tubes, both were transported to the Universitat Autònoma de Barcelona. Lymphocyte isolation was performed, from the sodium heparin tubes, using lymphosep solution (Biowest, Nuaillé, France) following the manufacturer’s protocol. Once isolated, lymphocytes were diluted in 10.5 ml of RPMI 1640 (Biowest) supplemented with 15% of fetal bovine serum (FBS) (Life Technologies, Madrid, Spain). Then, lymphocyte suspension was split into seven different tubes containing 1.5 ml each one. Five of these tubes were used for the γ-H2AX assay and two for the apoptosis assay. DNA extraction was carried out from the EDTA tubes.

### 2.3 Irradiation conditions

For the γ-H2AX assay, four tubes were irradiated at 2 Gy and one tube was sham-irradiated. For the apoptosis assay, one tube was irradiated at 8 Gy and another tube was sham-irradiated. Lymphocyte suspensions were irradiated with a ^137^Cs-source (IBL437C, CIS Bio international, Gif-Sur-Yvette, France) located in a separated building in the Universitat Autònoma de Barcelona. After irradiation, lymphocytes were placed on ice during transportation for 10 min, to halt DNA repair. Once in the laboratory, lymphocytes were incubated for different time points at 37°C. For the γ-H2AX assay, the time points were 1, 2, 4 and 24 h and for the apoptosis assay 48 h.

### 2.4 γ-H2AX immunofluorescence staining and flow cytometry analysis

To detect the presence of γ-H2AX, first, cells were fixed in 2% paraformaldehyde (Sigma-Aldrich Quimica, Madrid, Spain) for 5 min at 4°C and then cells were centrifuged at 600 g for 5 min. After that, the supernatant was eliminated and ethanol 70% was added at least for 2 h at -20°C. Then, cells were centrifuged at 600 g for 5 min, next the supernatant was eliminated and cells were resuspended with 1xPBS and centrifuged again at 600 g for 5 min. Afterwards, 1xPBS-0.5% Tritonx100 (Sigma-Aldrich) was added for 10 min for cell permeabilization. Then, cells were washed with 1xPBS. A blocking treatment was conducted for 30 min with 1xPBS-0.1% Tween20 (Sigma-Aldrich) and 2% of FBS. Next, cells were centrifuged and after supernatant removal, they were incubated overnight at 4°C with the mouse monoclonal anti-γ-H2AX (Ser139) antibody (Abcam, Cambridge, UK) diluted 1:500 in 1XPBS-0.1% Tween20 and 2% FBS. After this first incubation, cells were washed twice with 1xPBS-0.1% Tween20 solution and a second incubation was done using a secondary anti-mouse antibody, labelled with FITC (Invitrogen, Thermo-Fisher, Eugene, OR, USA) diluted 1:400 in 1XPBS-0.1% Tween20 and 2% FBS. DNA staining was done with a solution of 1 µl of propidium iodide (PI) (1 mg/ml) and 1 µl of RNase (1 mg/ml) in 1 ml of 1xPBS.

Analysis of γ-H2AX was done using the Cytoflex flow cytometer (Beckman Coulter, Indianapolis, IN, USA). This cytometer has two active lasers of 488 nm and 638 nm. To measure the γ-H2AX intensity of fluorescence in suspension, FITC and PI were excited with the 488 nm laser to detect γ-H2AX and DNA content respectively. For each sample the fluorescence intensity of approximately 10.000 events was recorded. To evaluate γ-H2AX fluorescence, the geometric Mean Fluorescence Intensity of FITC (MFI-FITC) was obtained by using FlowJo vX.0.7 software (FlowJo, Ashland, OR, USA) [48].

### 2.5 Radio-induced apoptosis assay with Annexin V/PI

To perform the apoptosis assay, the Annexin-V-FLUOS-Staining Kit (Roche, Barcelona, Spain) was used according to manufacturer’s instructions. Apoptosis was evaluated in the CD8+ lymphocyte population. For this reason, the APC anti-human CD8+ mouse antibody, SK1 clone, (Biolegend, San Diego, CA, USA) was added to the kit mix. Viable, Early Apoptotic (EA) and Late Apoptotic (LA) cells were scored by the Cytoflex flow cytometer. The Annexin V positive cells were identified using the 638 nm laser and the PI positive cells using the 488 nm laser. For each sample approximately 50.000 events were counted. Results were then analyzed using the FlowJo vX.0.7 software.

To evaluate the radio-induced cell death, each irradiated sample was normalized by subtracting its own non-irradiated control. Live cells were defined as cells negative for Annexin V and PI. EA cells were defined as cells positive for Annexin V and negative for PI and LA cells were defined as cells positive for both Annexin V and PI.

### 2.6 DNA extraction, quantification and SNP genotyping

DNA was extracted from blood samples using the QIAamp DNA Blood Midi Kit (QIAGEN, Hilden, Germany) and quantified using the Qubit dsDNA BR Assay Kit (Invitrogen) following manufacturer’s instructions for both kits. All DNA samples were diluted to a final concentration between 30 and 50 ng/ul. DNA was transferred to an Open Array Plate (Thermo-Fisher) with 96 wells. A total of 27 SNPs in genes related to apoptosis selected from the literature were evaluated. Studied SNPs are located in the following genes: *ADRB2, BAX, BCL2, BIRC5, CASP8, FAS, FASLG, GSTM1, GSTP1, iASPP, LGALS3, MCL1, MDM2, PDCD5, PDE4B, RASSF1, TNF* and *TP53*. Specific studied positions can be seen in Supplementary material 1.

### 2.7 Statistical analysis

Data were analyzed by GraphPad Prism version 8.0.1 (GraphPad Software, Boston, MA, USA) setting as statistically significant a p value < 0.05. First, normality tests (Shapiro-Wilk test and Kolmogorov-Smirnov test) were performed to γ-H2AX analysis values. After that, Wilcoxon test between the different time points was performed. Apoptosis assay was analyzed with t-test between two groups corresponding to 24 h and 48 h post-irradiation. Correlation of individual values, either for γ-H2AX results or apoptosis rate, was tested also with Spearman test.

In the genotyping assay, potential prognostic variables, in this case SNPs genotype and covariates, such as clinical parameters, were evaluated to know their implication in cell death. First, two different apoptosis groups were established using the median as a cut-off. Genetic variants were assessed for association with apoptosis using R 4.2.1 software [49] and R package SNPassoc 2.1.0 [50]. First, Hardy-Weinberg equilibrium was explored for the 27 SNPs using the Chi-square test. Then, logistic regression was used to test for association under five genetic models for each SNP (codominant, dominant, recessive, overdominant and log-additive). Moreover, some covariates were added to analyze their implication in the model. These covariates were qualitative, such as type of tumour and nodule, presence of metastasis, stage of the tumour, smoking habit, dyslipidemia, chemotherapy, hormonotherapy and monoclonal antibody therapy and others were quantitative, such as age, total RT dose and boost dose. Bonferroni correction was applied to adjust for multiple testing.

## 3. Results

### 3.1 γ-H2AX assay

Descriptive statistics of the results of γ-H2AX analysis are shown in Table 1. There was a wide range of the observed values for the MFI-FITC; in the sham-irradiated samples and the irradiated samples evaluated at different post-irradiation times. To figure out if some values were significant outliers, the ROUT test was applied, using as a Q value the recommended 1%. Out of the 300 measurements, five were considered as outliers (two in the sham-irradiated samples, one in the analysis performed 2 h post-irradiation and two in the 4 h post-irradiation time). Once these outlier values were removed, normality was evaluated using the Kolmogorov-Smirnov test. Only the MFI-FITC values 24 h post-irradiation agrees with the normal distribution. For this reason, further comparisons were done using non-parametric tests.

**Table 1.** Descriptive statistics of the geometric Mean Fluorescence Intensity of FITC (MFI-FITC) values after γ-H2AX assay. a) Crude data; b) ROUT correction.

| a) | Time after irradiation |  |  |  |  |
| --- | --- | --- | --- | --- | --- |
|  | NI | 1 h | 2 h | 4 h | 24 h |
|  | N = 60 | N = 60 | N = 60 | N = 60 | N = 60 |
| Maximum | 195470 | 321368 | 330946 | 324173 | 214257 |
| Minimum | 48474 | 63915 | 58295 | 68420 | 42324 |
| Median | 72820 | 138462 | 129690 | 118745 | 107953 |
| SD | 29524 | 53029 | 57117 | 48682 | 37367 |
| Mean | 81411 | 149122 | 145154 | 131840 | 108569 |
| SEM | 3812 | 6846 | 7374 | 6285 | 4824 |
| CV | 36.27% | 35.56% | 39.35% | 36.92% | 34.42% |

| b) | Time after irradiation |  |  |  |  |
| --- | --- | --- | --- | --- | --- |
|  | NI | 1 h | 2 h | 4 h | 24 h |

|  | N = 58 | N = 60 | N = 59 | N = 58 | N = 60 |
| --- | --- | --- | --- | --- | --- |
| Maximum | 136750 | 321368 | 304116 | 234972 | 214257 |
| Minimum | 48474 | 63915 | 58295 | 68420 | 42324 |
| Median | 72494 | 138462 | 128748 | 117529 | 107953 |
| SD | 22672 | 53029 | 52090 | 37808 | 37367 |
| Mean | 77865 | 149122 | 142005 | 126119 | 108569 |
| SEM | 2977 | 6846 | 6781 | 4964 | 4824 |
| CV | 29.12% | 35.56% | 36.68% | 29.98% | 34.42% |

NI, non-irradiated; N, number of patients; SD, standard deviation; SEM, standard error of the mean; CV, coefficient of variation.

The MFI-FITC was significantly different between each post-irradiation time point, except between 1 and 2 h post-irradiation (Fig 1). The basal or non-irradiated (NI) value of γ-H2AX intensity median level was always significantly lower than those observed after irradiation, even after 24 h of repair. Individual MFI-FITC values observed in the sham-irradiated samples were positively correlated to those reported after 1, 2, 4 and 24 h post irradiation. The R values were 0.38 (p = 0.003), 0.44 (p = 0.001), 0.56 (p < 0.0001) and 0.57 (p < 0.0001), respectively.

**Figure 1.**
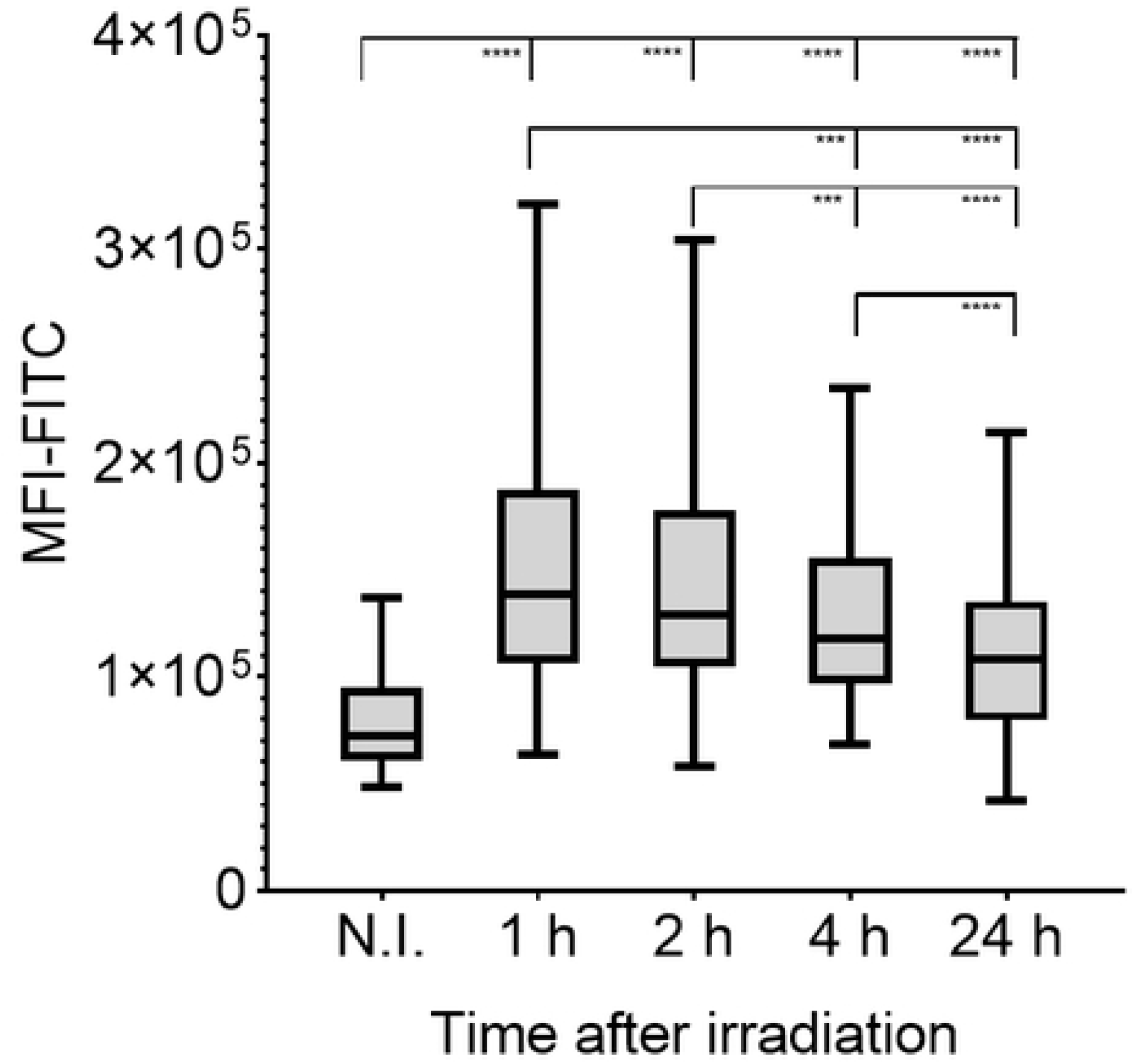
Box-plot with mean geometric fluorescence intensity of FITC (MFI-FITC). Values for each evaluated condition, non-irradiated (NI) and irradiated at 2 Gy, analyzed 1, 2, 4 and 24 h after irradiation. (****p < 0.0001, ***p < 0.001).

### 3.2 Apoptosis assay

For each individual, the EA and LA (or necrotic) percentages from CD8^+^ observed after 8 Gy irradiation were considered jointly after subtracting the respective values observed in the sham-irradiated samples (Table 2); to avoid outliers, negative results were removed. As can be seen in Fig 2, the percentage of radio-induced apoptosis at 48 h post irradiation was statistically higher than that observed at 24 h.

**Figure 2.**
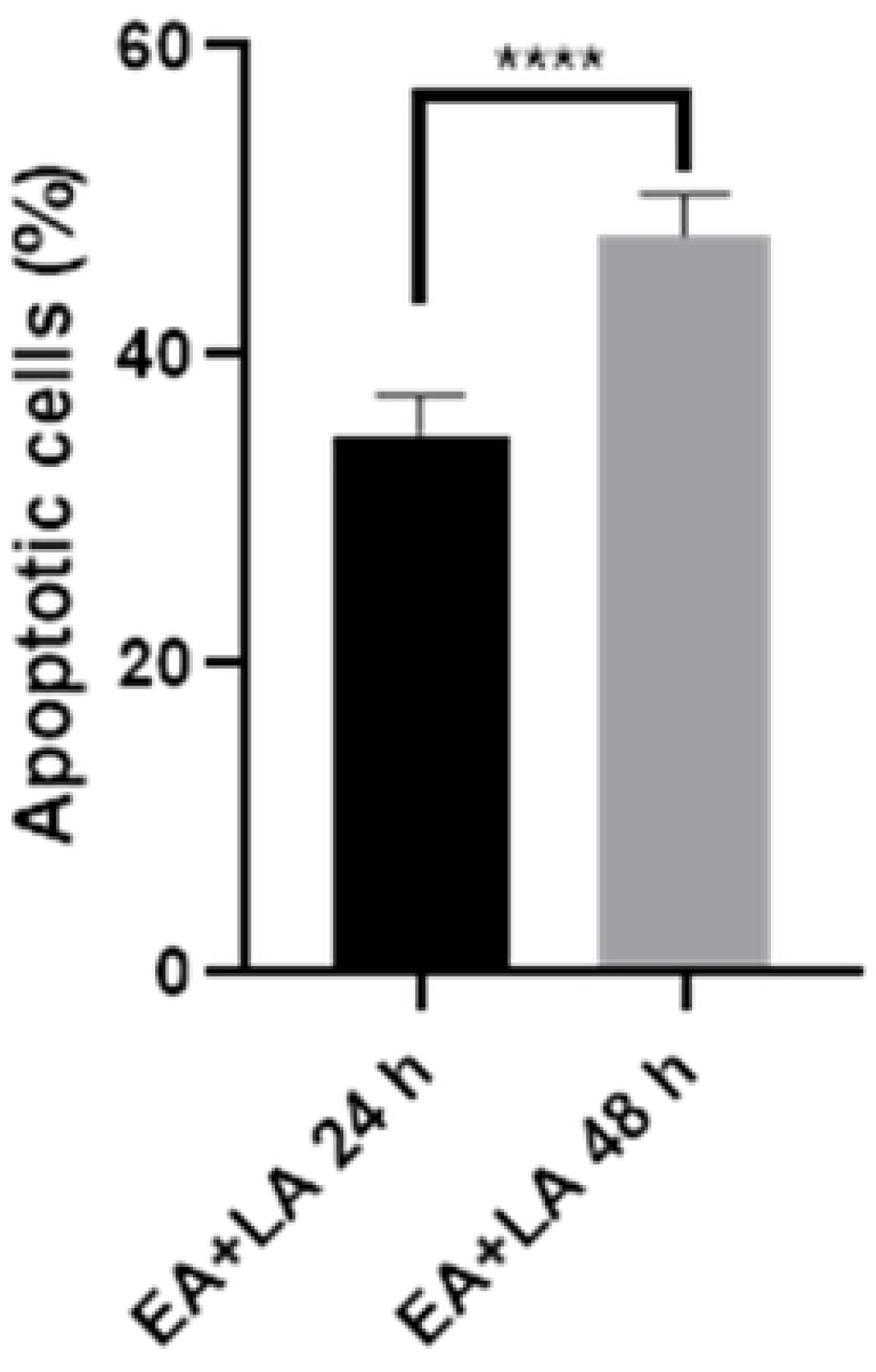
Percentage of radio-induced total apoptotic cells (Early and Late-apoptosis, EA+LA). Evaluation after 24 h and 48 h post irradiation at 8 Gy (**** p < 0.0001). Mean values and ± SEM values were used.

**Table 2.**
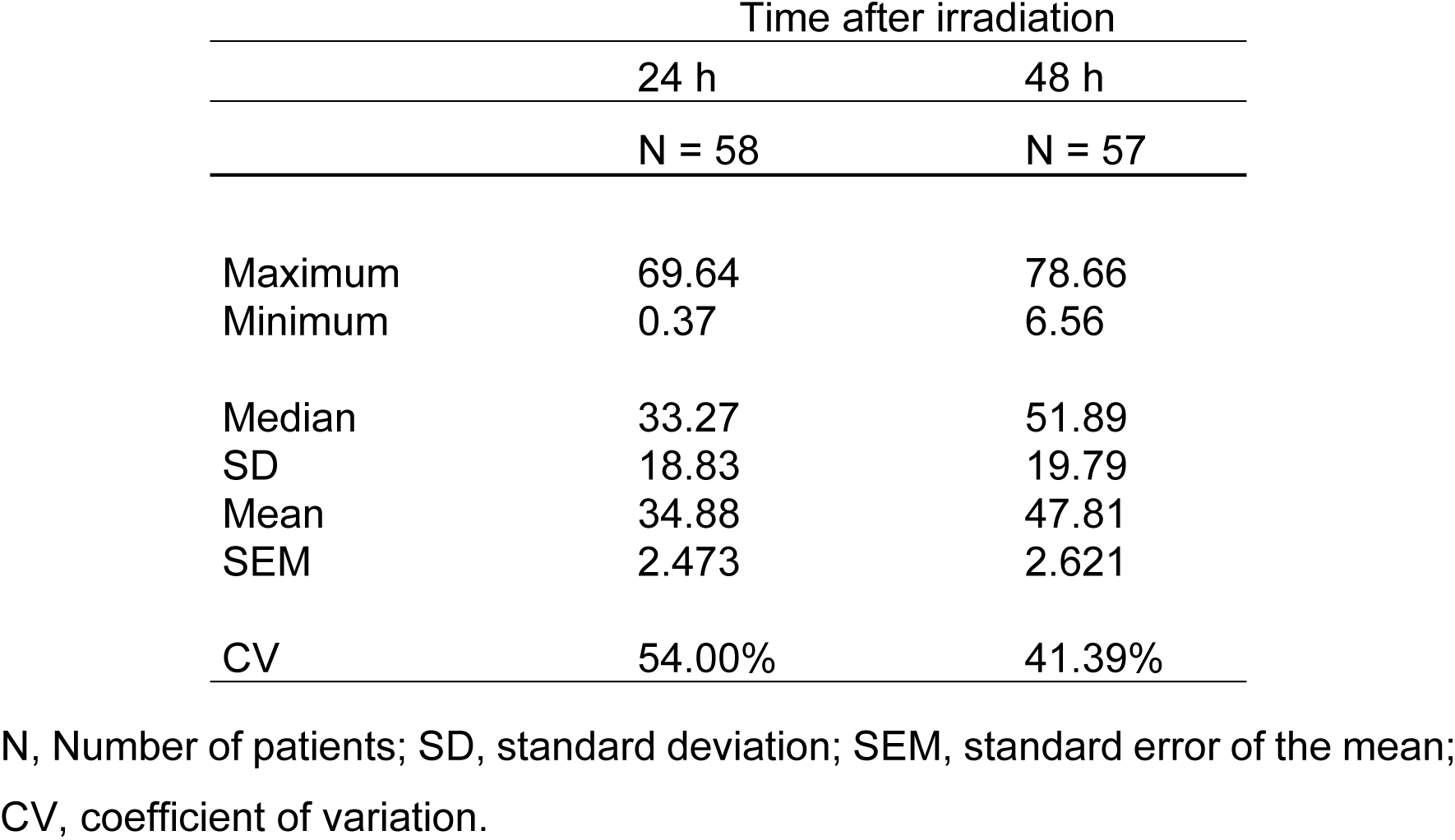
Descriptive statistics of the radio-induced apoptosis rate.

|  | Time after irradiation |  |
| --- | --- | --- |
|  | 24 h | 48 h |
|  | N = 58 | N = 57 |
| Maximum | 69.64 | 78.66 |
| Minimum | 0.37 | 6.56 |
| Median | 33.27 | 51.89 |
| SD | 18.83 | 19.79 |
| Mean | 34.88 | 47.81 |
| SEM | 2.473 | 2.621 |
| CV | 54.00% | 41.39% |

N, Number of patients; SD, standard deviation; SEM, standard error of the mean; CV, coefficient of variation.

To study deeply the reason of these differences, EA and LA at each time point were studied separately. In the irradiated samples and for both post-irradiation times 24 and 48 h, the percentages of EA and LA cells showed differences (Fig 3). After 24 h the percentage of EA cells was higher than that after 48 h (11.240 ± 7.183 vs. 8.635 ± 7.182, p = 0.04); and inversely after 48 h the percentage of LA cells was higher than after 24 h (43.84 ± 16.28 vs. 25.66 ± 14.92, p = 0.0001).

**Figure 3.**
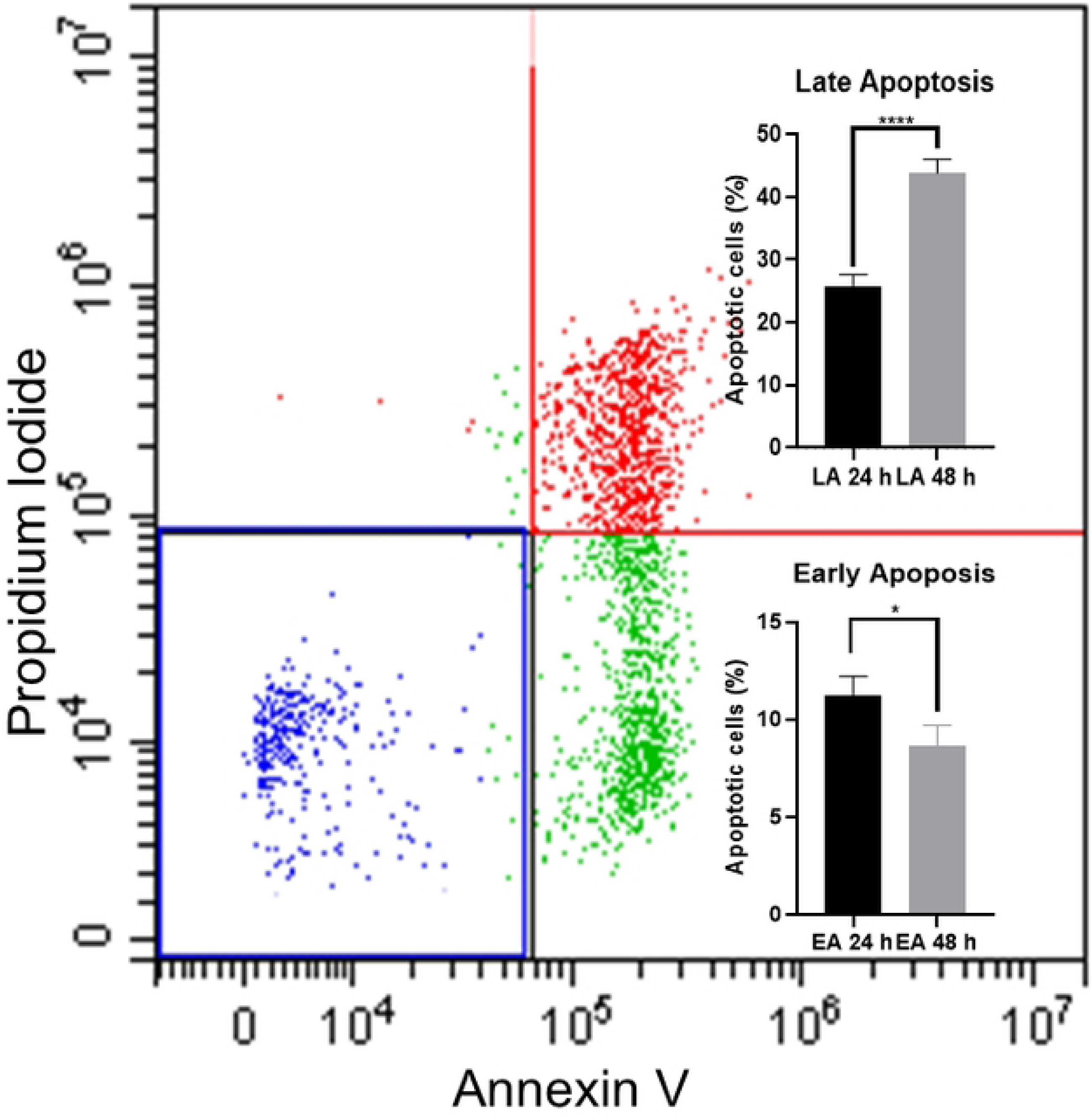
Radio-induced apoptosis measurement for a representative sample after 8 Gy. Live (Annexin V and Propidium Iodide negative) cells are presented in blue, early apoptotic cells (Annexin V positive and Propidium Iodide negative) presented in green and late apoptotic cells (Annexin V and Propidium Iodide positive) cells presented in red. Embedded figures show early (EA) and late apoptotic cells (LA) analyzed at 24 and 48 h after irradiation (* p<0.05, **** p<0.0001). Mean and ± SEM values were used.

For both 24 and 48 h post-irradiation times, a positive correlation was noticed between the individual values of EA and LA in sham-irradiated samples and the EA and LA values after 8 Gy irradiation (p < 0.05).

### 3.3 Correlation between γ-H2AX and apoptosis

The potential correlation between the apoptotic rate at 24 and 48 h and the γ-H2AX results was evaluated. After 8 Gy irradiation and after 24 h of incubation, individual level of apoptotic cells (considering together EA and LA) did not show any correlation with the observed MFI-FITC values at any post-irradiation time. However, after 48 h post-irradiation the apoptosis values showed a negative correlation with the basal values of MFI-FITC (R = -0.31, p = 0.023) and with radiation-induced MFI-FITC observed 24 h post-irradiation (R = -0.41, p = 0.002) (Fig 4).

**Figure 4.**
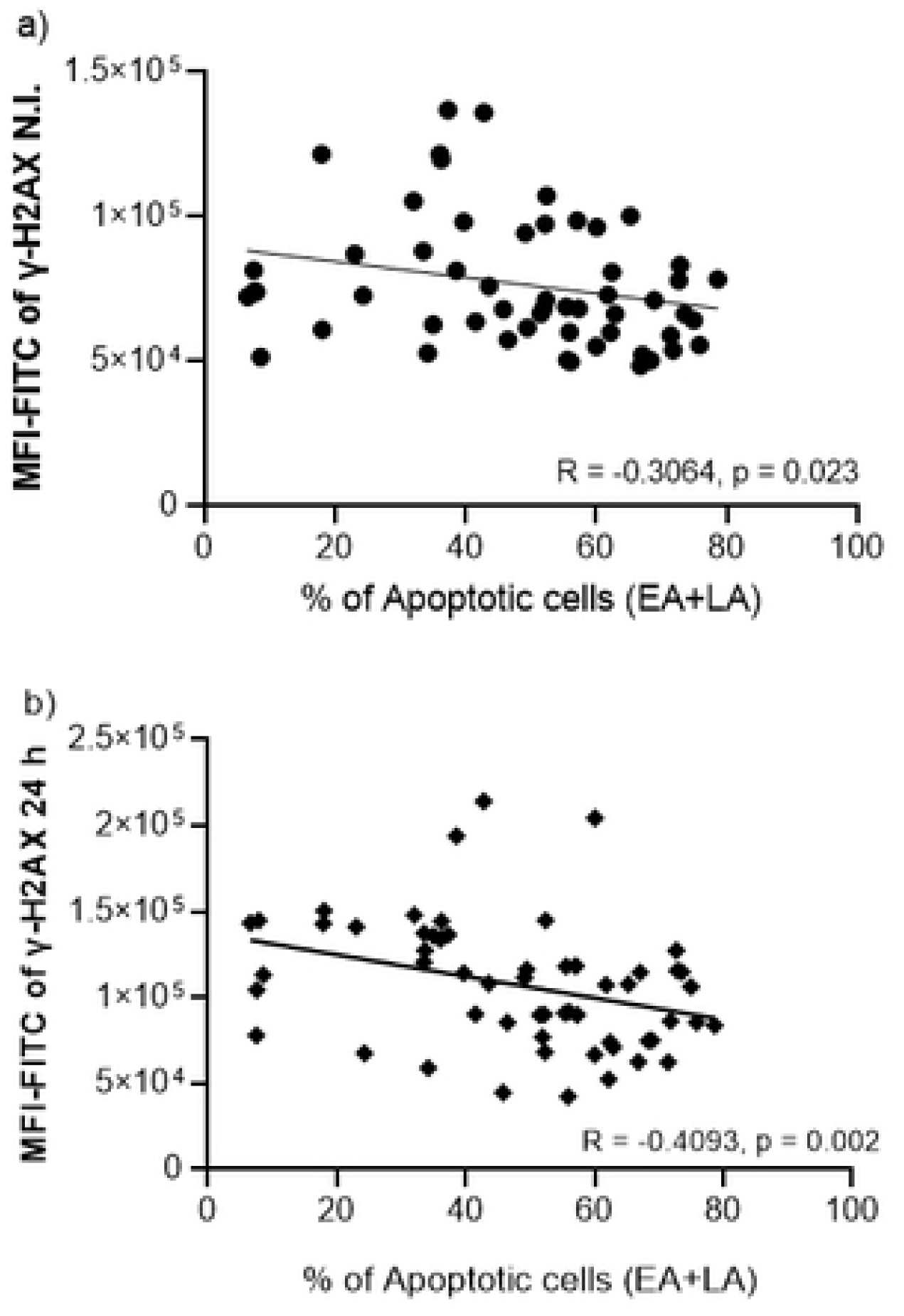
Correlation between the percentage of apoptotic cells and the geometric Means Fluorescence Intensity of FITC (MFI-FITC) of γ-H2AX. a) Apoptosis was measured after 48 h post-irradiation and MFI-FITC was measured in non-irradiated (N.I.) cells; and b) Apoptosis was measured after 48 h post-irradiation and MFI-FITC was measured after 24 h post-irradiation at 2 Gy.

### 3.4 SNP genotyping

The genotype distribution of all SNPs followed the Hardy-Weinberg equilibrium, except for *TNF* rs1799724 (p = 0.03). Out of the 27 SNPs evaluated, SNPassoc analysis found differences between the two groups of apoptosis rate in: *TP53* rs1625895 (C/C vs C/T, OR = 6.88, 95% confidence interval (CI) [1.71-27.75], p = 0.003) in codominant model, *FAS* rs1800682 (G/G vs G/A+A/A, OR = 0.18, 95% CI [0.05-0.63], p = 0.004) in dominant model, *BCL2L11* rs724710 (C/C vs C/T+T/T, OR = 0.32, 95% CI [0.11-0.94], p = 0.034) in dominant model and *MCL1* rs9803935 (G/G+G/T vs T/T OR = 0.27, 95%CI [0.07-0.96], p = 0.034) in recessive model (Table 3). After using Bonferroni correction and for α = 0.05, none of them were statistically significant. With α = 0.1, two statistically significant genes were obtained: *TP53* rs1625895 in codominant model and *FAS* rs1800682 in dominant model. Any covariate, neither age nor the total dose received during the treatment, for example, was statistically significant in our analysis.

**Table 3.** Association of selected polymorphisms with apoptosis.

| Polymorphism |  |  | Group of study |  |  | P value |  |  |  |  |
| --- | --- | --- | --- | --- | --- | --- | --- | --- | --- | --- |
| SNP | Gene | Genotype | Lower Apoptosis | Higher Apoptosis | Total | Codominant | Dominant | Recessive | Overdominant | Log AD |
| rs1625895 | TP53 | C/C | 27 | 17 | 44 | 0.003* | - | - | - | - |
|  |  | C/T | 3 | 13 | 16 |  |  |  |  |  |
|  |  | T/T | 0 | 0 | 0 |  |  |  |  |  |
| rs1800682 | FAS | G/G | 4 | 14 | 18 | 0.016 | 0.004* | 0.345 | 0.069 | 0.017 |
|  |  | A/G | 18 | 11 | 29 |  |  |  |  |  |
|  |  | A/A | 8 | 5 | 13 |  |  |  |  |  |
| rs724710 | BCL2L11 | C/C | 8 | 16 | 24 | 0.087 | 0.034 | 1 | 0.038 | 0.104 |
|  |  | C/T | 19 | 11 | 30 |  |  |  |  |  |
|  |  | T/T | 3 | 3 | 6 |  |  |  |  |  |
| rs9803935 | MCL1 | G/G | 8 | 7 | 15 | 0.06 | 0.766 | 0.034 | 0.038 | 0.272 |
|  |  | G/T | 11 | 19 | 30 |  |  |  |  |  |
|  |  | T/T | 11 | 4 | 15 |  |  |  |  |  |

SNP, Single Nucleotide Polymorphism; Log AD, Log-Additive
\*Statistically significant p value after Bonferroni correction ( $\alpha = 0.1$ ).

## 4. Discussion

Ionizing radiation is able to produce several types of DNA damage and all of them activate specific DNA repair mechanisms [51]. Among the distinct types of DNA damage, DSB, which occur in smaller amounts compared to other DNA damage types, are considered the ones with more biological impact. DSB are related to rearrangements of genomic material (inducing chromosomal aberrations) and to cell death [52,53]. The analysis of γ-H2AX allows to evaluate DSB induction as well as their repair kinetics. The γ-H2AX assay is widely used to study factors that can influence the DNA damage response and has been proposed to be used in clinical studies to detect radiosensitive patients [54,55]. In the present study, the highest value of γ-H2AX intensity was observed at 1 h post-irradiation, which agrees with other studies proving that the maximum levels of γ-H2AX occur between 0.5 and 1 h [56]. Notably, a big variability was observed in the results, with CVs higher than 29% even after outlier correction. Furthermore, the individual γ-H2AX values were considered at each evaluated condition and a positive correlation was observed between the basal values and the ones obtained at each post-irradiation time point including 24 h irradiation. Since constitutive levels of γ-H2AX can be considered to show the capacity to solve endogenous DNA damage [57], it could be hypothesized that the individual differences are related to the DSB repair capacity [58]. Our results seem to indicate that those individuals less efficient in resolving endogenous DSBs are also those that show lower efficiency in resolving radiation-induced DSBs. Interestingly, in a previous study evaluating 94 healthy individuals, age, alcohol consumption and ethnicity were found to influence baseline γ-H2AX levels, whereas ethnicity and alcohol influenced repair kinetics [59]. In our study, all women belonged to the same ethnic group and alcohol consumption was not reported. When considering age, no correlation was seen with either initial γ-H2AX levels or repair kinetics. This shows that even a homogeneous group can present differences in DNA damage induction or repair.

After exposure to ionizing radiation, in case of severe or irreparable DNA and to prevent cells with mutations from progressing or producing genomic instability, cells can trigger cell death [35]. Currently, many forms of programmed cell death are described: intrinsic or extrinsic apoptosis, necrosis, ferroptosis, piroptosis and mitotic catastrophe, among others [60]. In G0 lymphocytes, ionizing radiation mainly promotes intrinsic apoptotic pathway [61]. Radiation induced damage in the intracellular structures like DNA, activates the intrinsic pathway releasing cytochrome C from mitochondria. In addition, ionizing radiation can also promote the extrinsic apoptosis, which is started by death receptors present in the cell membrane. In G0 cells, another conceivable way to produce apoptosis would be due to the membrane stress. In this case, radiation induced reactive oxygen species oxidate cell membrane lipids triggering this cell death pathway. In the present study, cell death was evaluated 24 and 48 h post-irradiation and data presented large variability, with CVs higher than 41%. When considering radiation-induced apoptosis by subtracting the corresponding controls, the percentage of total apoptotic cells (EA+LA) was higher 48 h after irradiation. Our results confirm that apoptosis can be better evaluated after 48 h post-irradiation rather than after 24 h [37,62], which may be because the EA cells at 24 h become LA cells at 48 h, increasing the percentage and in consequence the statistical power.

When the two biomarkers of radiation sensitivity, γ-H2AX and apoptosis, were correlated, the results indicated a significant negative correlation between the percentage of apoptotic cells, induced after 8 Gy and evaluated 48 h post-irradiation and the γ-H2AX values observed in the non-irradiated samples and in the samples irradiated at 2 Gy and analyzed 24 h later. In a previous study with 16 breast cancer patients, Chua et al. [39] found a negative, although non-significant, correlation between the mean number of γ-H2AX/53BP1 foci observed 24 h post-irradiation at 4 Gy and the mean apoptosis rate 48 h post-irradiation at 8 Gy. Later the same group [63], evaluating 172 nasopharynx cancer patients, also saw a significant inverse association between residual DSB and percentage of apoptosis. These authors suggested that the negative correlation was due to a subgroup of patients with some intrinsic genomic or epigenomic factors that result in a blunting of the DNA damage response. Other authors have also observed this negative correlation, evaluating breast cancer patients before the start and at 1 and 2 weeks of RT [64]. Our results reinforce the existence of this negative correlation between apoptosis and background and residual levels of γ-H2AX. The negative correlation by using “ex vivo” irradiated lymphocytes suggests that those individuals that are more efficient at removing damaged cells through apoptosis are those that show lower levels of background or residual levels of DNA damage.

To assess whether the variability observed in apoptosis rate was related to genetic polymorphisms, we performed a SNP analysis. Patients were divided into two groups depending on their percentage of total radio-induced apoptotic cells. The analysis showed that two SNPs were differentially represented in the two groups. The first SNP was rs1625895 in the gene coding for TP53, which is activated in the presence of DNA damage and it also engages in the intrinsic apoptosis pathway. This transcription factor induces the production of pro-apoptotic proteins of the Bcl-2 family, which allow the release of cytochrome C and the subsequent activation of Caspase-9, starting the cell death process itself [65]. In both groups of patients, the most common genotype was CC, as described in the general population [66]. The positive association of this SNP with the radio-induced apoptosis rate followed a codominant model and CT genotype was more present in patients that showed elevated percentages of radio-induced apoptotic cells. Accordingly, the T allele has been observed in cells having a greater ability to enter apoptosis [67].

The second SNP that was differentially represented in the two groups was rs1800896 in the gene coding for FAS, which is related to the extrinsic pathway of apoptosis. It is known that radiation causes an increase in the production of ligands, such as FasL and TNF-alfa, which bind to their cell membrane receptors to form the Death Inducing Cell Signalling Complex. This complex activates the Caspase-8, which triggers apoptosis [60]. For this SNP, our patients mostly presented the AG genotype, which agrees with what is expected from the general population [66]. In our study, this SNP showed a negative association with the radio-induced apoptosis rate in a dominant genetic model. Previous studies have reported that the G allele is associated with a lower apoptosis level but also with a higher risk to develop breast cancer [68,69]. Contrary to expectations, our results shown that GG genotype is more present to the group with higher radio-induced apoptosis. A possible explanation for this contradictory result could be that this SNP is also associated with an increase of risk to develop breast cancer.

Since our cohort is only composed of breast cancer patients, this can mask the statistical output.

Based on this result, we checked whether those patients with the higher apoptosis rate who presented the CT genotype in rs1625895 are the same as those with GG and AG genotypes for the rs1800682. Out of 13 patients with CT for the rs1625895, 10 patients have the GG (5 patients) or AG (5 patients) genotype of rs1800682. This indicates that despite having genotypes (GG or AG) related to low levels of extrinsic apoptosis, the CT genotype, related to higher levels of intrinsic apoptosis, seems to have greater weight. These results support the idea that the intrinsic pathway, due to DNA damage, appears to play a more key role in radiation-induced apoptosis than the extrinsic pathway [61].

In conclusion, the results here presented demonstrate that breast cancer patients with less efficiency in removing damaged cells, have more background and residual levels of DNA damage and that this variability could be explained by individual genetic variations. In this sense, further studies will be carried out to test whether these and other biomarkers are associated with adverse RT effects in these patients.

## Acknowledgement

We want to thank Nil Campderrós for his laboratory and analysis support, Jessica Martinez-Cortes for her laboratory work and Karen Mieles, Yasmina Castillo, Sandra Clotet, Gerard Rodríguez for their clinical work.

## Authors contribution

**Conceptualization:** Monica Pujol-Canadell, Miquel Macià, Evelyn Martínez Pérez, Ferran Guedea, Victor González-Rumayor, Gemma Armengol, Joan Francesc Barquinero

**Data curation:** Ángela Solana-Peña, Juan-Sebastián López

**Formal analysis:** Ángela Solana-Peña, Monica Pujol-Canadell, Gemma Armengol, Joan Francesc Barquinero

**Funding Acquisition:** Ferran Guedea, Victor González-Rumayor, Joan Francesc Barquinero

**Investigation:** Ángela Solana-Peña, Monica Pujol-Canadell, Juan-Sebastián López

**Methodology:** Monica Pujol-Canadell

**Project administration:** Monica Pujol-Canadell, Gemma Armengol, Joan Francesc Barquinero

**Resources:** Miquel Macià, Evelyn Martínez Pérez, Isabel Linares, Milica Stefanovic, Héctor Pérez-Montero, Javier González-Viguera, Marina Arangüena Peñacoba, Ferran Guedea.

**Software:** Ángela Solana-Peña

**Supervision:** Gemma Armengol, Joan Francesc Barquinero

**Visualization:** Ángela Solana-Peña, Monica Pujol-Canadell, Montse Ventura, Gisela de Miguel-Garcia, Ferran Guedea, Nadina Erill, Victor González-Rumayor, Gemma Armengol, Joan Francesc Barquinero,

**Writing-original draft:** Ángela Solana-Peña, Joan Francesc Barquinero

**Writing-review and editing:** Ángela Solana-Peña, Monica Pujol-Canadell, Gemma Armengol, Joan Francesc Barquinero

## Funding

This work was supported by the Plan Estatal de Investigación Científica, Técnica y de Innovación 2021-2023, in the frame of Plan de Recuperación, Transformación y Resiliencia, (Proyectos de Colaboración Público-Privada, CPP2021-008368). Ministerio de Ciencia e Innovación.

## Data Availability

Research data are stored in the Catalan Open Research Area (CORA), https://doi.org/10.34810/data18

## Notes

### Competing Interest Statement

The authors have declared no competing interest.

## References

1. American Cancer Society. Information and resources about cancer: breast, colon, lung, prostate, skin [Internet]. 2024 [cited 2024 Sep 23]. Available from: https://www.cancer.org

2. Deloch L, Derer A, Hartmann J, Frey B, Fietkau R, Gaipl US. Modern radiotherapy concepts and the impact of radiation on immune activation. Front Oncol. 2016;6: 1–16. doi:10.3389/fonc.2016.00141

3. Majeed H, Gupta V. Adverse effects of radiation therapy [Internet]. In: StatPearls. StatPearls Publishing; 2023 [cited 2024 Dec 17]. Available from: https://www.ncbi.nlm.nih.gov/books/NBK563259/

4. Yarnold J. Changes in radiotherapy fractionation-breast cancer. Br J Radiol. 2019;92.

5. LaRiviere MJ, Vapiwala N. Radiation Therapy. Penn Clinical Manual of Urology, 3rd ed. Philadeohia: Elsevier; 2022.p.704–734.e5. doi:10.1016/B978-0-323-77575-5.00028-9

6. Greve B, Bö Lling T, Amler S, Rö Ssler U, Gomolka M, Mayer C, et al. Evaluation of Different Biomarkers to Predict Individual Radiosensitivity in an Inter-Laboratory Comparison-Lessons for Future Studies. 2012doi:10.1371/journal.pone.0047185

7. Barnett GC, West CML, Dunning AM, Elliott RM, Coles CE, Pharoah PDP, et al. Normal tissue reactions to radiotherapy: towards tailoring treatment dose by genotype. Nat Rev Cancer. 2009;9: 134. doi:10.1038/NRC2587

8. Hsu TC. Genetic instability in the human population: a working hypothesis. Hereditas. 1983;98:1–9. doi:10.1111/j.1601-5223.1983.tb00760.x

9. Bentzen SM, Overgaard J. Patient-to-patient variability in the expression of radiation-induced normal tissue injury. Semin Radiat Oncol. 1994;4: 68–80. doi:10.1016/S1053-4296(05)80034-7

10. Constine LS, Olch AJ, Jackson A, Hua C-H, Ronckers CM, Milano MT, et al. Pediatric Normal Tissue Effects in the Clinic (PENTEC): An International Collaboration to Assess Normal Tissue Radiation Dose-Volume-Response Relationships for Children With Cancer. Int J Radiat Oncol Biol Phys. 2021; 119:316–20. doi:10.1016/J.IJROBP.2020.10.040

11. Turesson I, Nyman J, Holmberg E, Oden A. Prognostic factors for acute and late skin reactions in radiotheraphy patients. Int J Radiat Oncol Biol Phys. 1996;36: 1065–1075. doi:10.1016/S0360-3016(96)00426-9

12. Andreassen CN, Alsner J, Overgaard J. Does variability in normal tissue reactions after radiotherapy have a genetic basis - Where and how to look for it? Radiotherapy and Oncology. 2002;64: 131–140. doi:10.1016/S0167-8140(02)00154-8

13. Gomolka M, Blyth B, Bourguignon M, Badie C, Schmitz A, Talbot C, et al. Potential screening assays for individual radiation sensitivity and susceptibility and their current validation state. Int J Radiat Biol. 2020;96: 280–296. doi:10.1080/09553002.2019.1642544

14. Kolnohuz A, Ebrahimpour L, Yolchuyeva S, Manem VSK. Gene expression signature predicts radiation sensitivity in cell lines using the integral of dose-response curve. BMC Cancer. 2024;24. doi:10.1186/S12885-023-11634-3

15. Naderi E, Aguado-Barrera ME, Schack LMH, Dorling L, Rattay T, Fachal L, et al. Large-scale meta-genome-wide association study reveals common genetic factors linked to radiation-induced acute toxicities across cancer types. JNCI Cancer Spectr. 2023;7. doi:10.1093/JNCICS/PKAD088

16. Adams G, Martin OA, Roos DE, Lobachevsky PN, Potter AE, Mbbs Bm, et al. Enhanced intrinsic radiosensitivity after treatment with stereotactic radiosurgery for an acoustic neuroma. Radiother Oncol. 2012;103: 410–414. doi:10.1016/j.radonc.2012.03.011

17. González JE, Lee M, Barquinero JF, Valente M, Roch-Lefèvre S, García. O. Quantitative image analysis of gamma-H2AX foci induced by ionizing radiation applying open source programs. Anal Quant Cytol Histol. 2012;34: 66–71.

18. Schuler N, Palm J, Kaiser M, Betten D, Furtwä Ngler R, Rü Be C, et al. DNA-Damage Foci to Detect and Characterize DNA Repair Alterations in Children Treated for Pediatric Malignancies. PLoS One. 2014;17;9:3 doi: 10.1371/journal.pone.0091319.

19. Lobachevsky P, Leong T, Daly P, Smith J, Best N, Thompson ER, et al. Compromized DNA repair as a basis for identification of cancer radiotherapy patients with extreme radiosensitivity. Cancer Lett. 2016;383: 212–219. doi:10.1016/j.canlet.2016.09.010

20. Li P, Cheng-Run Du †, Xu W-C, Shi Z-L, Zhang Q, Li Z-B, et al. Correlation of dynamic changes in γ-H2AX expression in peripheral blood lymphocytes from head and neck cancer patients with radiation-induced oral mucositis. Radiation Oncology. 2013;8: 1. doi:10.1186/1748-717X-8-155

21. Van Oorschot B, Hovingh S, Dekker A, Stalpers LJ, Franken NAP. Predicting Radiosensitivity with Gamma-H2AX Foci Assay after Single High-Dose-Rate and Pulsed Dose-Rate Ionizing Irradiation. Radiat Res. 2016;185: 190–198. doi:10.1667/RR14098.1

22. Goutham HV, Mumbrekar KD, Vadhiraja BM, Fernandes DJ, Sharan K, Kanive Parashiva G, et al. DNA double-strand break analysis by γ-H2AX foci: a useful method for determining the overreactors to radiation-induced acute reactions among head-and-neck cancer patients. Int J Radiat Oncol Biol Phys. 2012;84. doi:10.1016/J.IJROBP.2012.06.041

23. Vinnikov V, Hande MP, Wilkins R, Wojcik A, Zubizarreta E, Belyakov O. Prediction of the Acute or Late Radiation Toxicity Effects in Radiotherapy Patients Using Ex Vivo Induced Biodosimetric Markers: A Review. J Pers Med. 2020;10: 1–36. doi:10.3390/JPM10040285

24. Matsubara S, Saito F, Suda T, Fuibayashi H, Shibuya H, Horiuchi J, et al. Radiation injury in a patient with unusually high sensitivity to radiation. Acta Oncol. 1988;27: 67–71. doi:10.3109/02841868809090321

25. Greulich-Bode KM, Zimmermann F, Muller W-U, Pakisch B, Molls M, Wurschmidt F. Clinical, molecular- and cytogenetic analysis of a case of severe radio-sensitivity. Curr Genomics. 2012;13: 426–432. doi:10.2174/138920212802510475

26. Fahrig A, Koch T, Lenhart M, Rieckmann P, Fietkau R, Distel L, et al. Lethal outcome after pelvic salvage radiotherapy in a patient with prostate cancer due to increased radiosensitivity: Case report and literature review. Strahlenther Onkol. 2018;194: 60–66. doi:10.1007/S00066-017-1207-9

27. Dunst J, Gebhart E, Neubauer S. Can an extremely elevated radiosensitivity in patients be recognized by the in-vitro testing of lymphocytes?. Strahlenther Onkol.1995;171:581–586.

28. Barber JBP, Burrill W, Spreadborough AR, Levine E, Warren C, Kiltie AE, et al. Relationship between in vitro chromosomal radiosensitivity of peripheral blood lymphocytes and the expression of normal tissue damage following radiotherapy for breast cancer. Radiotherapy and Oncology. 2000;55: 179–186. doi:10.1016/S0167-8140(99)00158-9

29. Brzozowska K, Pinkawa M, Eble MJ, Müller WU, Wojcik A, Kriehuber R, et al. In vivo versus in vitro individual radiosensitivity analysed in healthy donors and in prostate cancer patients with and without severe side effects after radiotherapy. Int J Radiat Biol. 2012;88: 405–413. doi:10.3109/09553002.2012.666002

30. Kondrashova T V., Ivanova TI, Katsalap SN. Chromosome aberrations in cultured peripheral lymphocytes from persons with elevated skin radiosensitivity. Environ Health Perspect. 1997;105 Suppl 6: 1437–1439. doi:10.1289/EHP.97105S61437

31. Borgmann K, Röper B, El-Awady RA, Brackrock S, Bigalke M, Dörk T, et al. Indicators of late normal tissue response after radiotherapy for head and neck cancer: Fibroblasts, lymphocytes, genetics, DNA repair, and chromosome aberrations. Radiotherapy and Oncology. 2002;64: 141–152. doi:10.1016/S0167-8140(02)00167-6

32. Borgmann K, Hoeller U, Nowack S, Bernhard M, Röper B, Brackrock S, et al. Individual radiosensitivity measured with lymphocytes may predict the risk of acute reaction after radiotherapy. Int J Radiat Oncol Biol Phys. 2008;71: 256–264. doi:10.1016/J.IJROBP.2008.01.007

33. Lee TK, Allison RR, O’Brien KF, Johnke RM, Christie KI, Naves JL, et al. Lymphocyte radiosensitivity correlated with pelvic radiotherapy morbidity. Int J Radiat Oncol Biol Phys. 2003;57: 222–229. doi:10.1016/S0360-3016(03)00411-5

34. Widel M, Jedrus S, Lukaszczyk B, Raczek-Zwierzycka K, Swierniak A. Radiation-induced micronucleus frequency in peripheral blood lymphocytes is correlated with normal tissue damage in patients with cervical carcinoma undergoing radiotherapy. Radiat Res. 2003;159: 713–721. doi: 10.1667/0033-7587(2003)159[0713:rmfipb]2.0.co;2

35. Biechonski S, Olender L, Zipin-Roitman A, Yassin M, Aqaqe N, Marcu-Malina V, et al. Attenuated DNA damage responses and increased apoptosis characterize human hematopoietic stem cells exposed to irradiation. Sci Rep. 2018;8: 1–13. doi:10.1038/s41598-018-24440-w

36. Ozsahin M, Crompton NEA, Gourgou S, Kramar A, Li L, Shi YQ, et al. CD4 and CD8 T-lymphocyte apoptosis can predict radiation-induced late toxicity: A prospective study in 399 patients. Clinical Cancer Research. 2005;11: 7426– 7433. doi:10.1158/1078-0432.CCR-04-2634

37. Azria D, Riou O, Castan F, Nguyen TD, Peignaux K, Lemanski C, et al. Radiation-induced CD8 T-lymphocyte Apoptosis as a Predictor of Breast Fibrosis After Radiotherapy: Results of the Prospective Multicenter French Trial. EBioMedicine. 2015;2: 1965–1973. doi:10.1016/j.ebiom.2015.10.024

38. Schnarr K, Boreham D, Sathya J, Julian J, Dayes IS. Radiation-Induced Lymphocyte Apoptosis to Predict Radiation Therapy Late Toxicity in Prostate Cancer Patients. Int J Radiat Oncol Biol Phys. 2009;74: 1424–1430. doi:10.1016/j.ijrobp.2008.10.039

39. Chua MLK, Horn S, Somaiah N, Davies S, Gothard L, A’Hern R, et al. DNA double-strand break repair and induction of apoptosis in ex vivo irradiated blood lymphocytes in relation to late normal tissue reactions following breast radiotherapy. Radiat Environ Biophys. 2014;53: 355–364. doi:10.1007/S00411-014-0531-Z/FIGURES/5

40. Cavalcante GC, Schaan AP, Cabral GF, Santana-Da-Silva MN, Pinto P, Vidal AF, et al. A cell’s fate: An overview of the molecular biology and genetics of apoptosis. Int J Mol Sci. 2019;20. doi:10.3390/ijms20174133

41. Mališić E, Petrović N, Brengues M, Azria D, Matić IZ, Srbljak Ćuk I, et al. Association of polymorphisms in TGFB1, XRCC1, XRCC3 genes and CD8 T-lymphocyte apoptosis with adverse effect of radiotherapy for prostate cancer. Sci Rep. 2022;12. doi:10.1038/S41598-022-25328-6

42. Dai YH, Wang YF, Shen PC, Lo CH, Yang JF, Lin CS, et al. Radiosensitivity index emerges as a potential biomarker for combined radiotherapy and immunotherapy. NPJ Genom Med. 2021;6. doi:10.1038/S41525-021-00200-0

43. Alsbeih G, El-Sebaie M, Al-Rajhi N, Al-Harbi N, Al-Hadyan K, Al-Qahtani S, et al. Among 45 variants in 11 genes, HDM2 promoter polymorphisms emerge as new candidate biomarker associated with radiation toxicity. 3 Biotech. 2014;4: 137–148. doi:10.1007/S13205-013-0135-3

44. Andreassen CN, Alsner J, Overgaard M, Sørensen FB, Overgaard J. Risk of radiation-induced subcutaneous fibrosis in relation to single nucleotide polymorphisms in TGFB1, SOD2, XRCC1, XRCC3, APEX and ATM--a study based on DNA from formalin fixed paraffin embedded tissue samples. Int J Radiat Biol. 2006;82: 577–586. doi:10.1080/09553000600876637

45. Barnett GC, Coles CE, Elliott RM, Baynes C, Luccarini C, Conroy D, et al. Independent validation of genes and polymorphisms reported to be associated with radiation toxicity: a prospective analysis study. Lancet Oncol. 2012;13: 65–77. doi:10.1016/S1470-2045(11)70302-3

46. Lumniczky K, Impens N, Armengol G, Candéias S, Georgakilas AG, Hornhardt S, et al. Low dose ionizing radiation effects on the immune system. Environ Int. 2021;149. doi:10.1016/J.ENVINT.2020.106212

47. Tam A, Mercier BD, Thomas RM, Tizpa E, Wong IG, Shi J, et al. Moving the Needle Forward in Genomically-Guided Precision Radiation Treatment. Cancers (Basel). 2023;15: 5314. doi:10.3390/CANCERS15225314

48. Wanotayan R, Chousangsuntorn K, Petisiwaveth P, Anuttra T, Lertchanyaphan W, Jaikuna T, et al. A deep learning model (FociRad) for automated detection of γ-H2AX foci and radiation dose estimation. Sci Rep. 2022;12. doi:10.1038/S41598-022-09180-2

49. R Core team (2023). R: A Language and Environment for Statistical Computing, https://www.r-project.org/

50. González JR, Armengol L, Solé X, Guinó E, Mercader JM, Estivill X, et al. SNPassoc: an R package to perform whole genome association studies. Bioinformatics. 2007;23: 644–645. doi:10.1093/BIOINFORMATICS/BTM025

51. Nikjoo H, Rahmanian S, Taleei R. Modelling DNA damage-repair and beyond. Prog Biophys Mol Biol. 2024;190: 1–18. doi:10.1016/J.PBIOMOLBIO.2024.05.002

52. Reisz JA, Bansal N, Qian J, Zhao W, Furdui CM. Effects of ionizing radiation on biological molecules--mechanisms of damage and emerging methods of detection. Antioxid Redox Signal. 2014;21: 260–292. doi:10.1089/ARS.2013.5489

53. Cheng B, Pan W, Xing Y, Xiao Y, Chen J, Xu Z. Recent advances in DDR (DNA damage response) inhibitors for cancer therapy. Eur J Med Chem. 2022;230. doi:10.1016/J.EJMECH.2022.114109

54. Dickey JS, Redon CE, Nakamura AJ, Baird BJ, Sedelnikova OA, Bonner WM. H2AX: functional roles and potential applications. Chromosoma. 2009;118: 683–692. doi:10.1007/S00412-009-0234-4

55. Barroso SI, Aguilera A. Detection of DNA Double-Strand Breaks by γ-H2AX Immunodetection. Methods Mol Biol. 2021;2153: 1–8. doi:10.1007/978-1-0716-0644-5_1

56. Rothkamm K, Horn S. γ-H2AX as protein biomarker for radiation exposure. Ann Ist Super Sanita. 2009; 45: 265–271.

57. Tanaka T, Halicka HD, Huang X, Traganos F, Darzynkiewicz Z. Constitutive histone H2AX phosphorylation and ATM activation, the reporters of DNA damage by endogenous oxidants. Cell Cycle. 2006;5: 1940–1945. doi:10.4161/CC.5.17.3191

58. Berthel E, Ferlazzo ML, Devic C, Bourguignon M, Foray N. What Does the History of Research on the Repair of DNA Double-Strand Breaks Tell Us?-A Comprehensive Review of Human Radiosensitivity. Int J Mol Sci. 2019;20. doi:10.3390/IJMS20215339

59. Sharma PM, Ponnaiya B, Taveras M, Shuryak I, Turner H, Brenner DJ. High Throughput Measurement of γH2AX DSB Repair Kinetics in a Healthy Human Population. PLoS One. 2015;10: e0121083. doi:10.1371/JOURNAL.PONE.0121083

60. Jiao Y, Cao F, Liu H. Radiation-induced Cell Death and Its Mechanisms. Health Phys. 2022;123: 376–386. doi:10.1097/HP.0000000000001601

61. Maier P, Hartmann L, Wenz F, Herskind C. Cellular Pathways in Response to Ionizing Radiation and Their Targetability for Tumor Radiosensitization. Int J Mol Sci. 2016;17. doi:10.3390/IJMS17010102

62. Foro P, Algara M, Lozano J, Rodriguez N, Sanz X, Torres E, et al. Relationship between radiation-induced apoptosis of T lymphocytes and chronic toxicity in patients with prostate cancer treated by radiation therapy: a prospective study. Int J Radiat Oncol Biol Phys. 2014;88: 1057–1063. doi:10.1016/J.IJROBP.2014.01.002

63. Chua KLM, Yeo ELL, Shihabudeen WA, Tan SH, Shwe TT, Ong EHW, et al. Intra-patient and inter-patient comparisons of DNA damage response biomarkers in Nasopharynx Cancer (NPC): Analysis of NCC0901 randomised controlled trial of induction chemotherapy in locally advanced NPC. BMC Cancer. 2018;18: 1–11. doi:10.1186/S12885-018-5005-2/FIGURES/5

64. Marková E, Somsedíková A, Vasilyev S, Pobijaková M, Lacková A, Lukačko P, et al. DNA repair foci and late apoptosis/necrosis in peripheral blood lymphocytes of breast cancer patients undergoing radiotherapy. Int J Radiat Biol. 2015;91: 934–945. doi:10.3109/09553002.2015.1101498

65. Yang S, Huang J, Liu P, Li J, Zhao S. Apoptosis-inducing factor (AIF) nuclear translocation mediated caspase-independent mechanism involves in X-ray-induced MCF-7 cell death. Int J Radiat Biol. 2017;93: 270–278. doi:10.1080/09553002.2016.1254833

66. Ensembl genome browser 112. [cited 5 Sep 2024]. Available: https://www.ensembl.org/index.html

67. Voropaeva EN, Voevoda MI, Pospelova TI, Maksimov VN. Prognostic impact of the TP53 rs1625895 polymorphism in DLBCL patients. Br J Haematol. 2015;169: 32–35. doi:10.1111/BJH.13237

68. Ji G, Gu A, Hu F, Wang S, Liang J, Xia Y, et al. Polymorphisms in cell death pathway genes are associated with altered sperm apoptosis and poor semen quality. Hum Reprod. 2009;24: 2439–2446. doi:10.1093/HUMREP/DEP223

69. Wang M, Wang Z, Wang XJ, Jin TB, Dai ZM, Kang HF, et al. Distinct role of the Fas rs1800682 and FasL rs763110 polymorphisms in determining the risk of breast cancer among Han Chinese females. Drug Des Devel Ther. 2016;10: 2359. doi:10.2147/DDDT.S111084

